# KSHV Reprograms Host RNA Splicing via FAM50A to Activate STAT3 and Drive Oncogenic Cellular Transformation

**DOI:** 10.1101/2025.03.17.643747

**Authors:** Shenyu Sun, Karla Paniagua, Ling Ding, Xian Wang, Yufei Huang, Mario A Flores, Shou-Jiang Gao

## Abstract

RNA alternative splicing is a fundamental cellular process implicated in cancer development. Kaposi’s sarcoma-associated herpesvirus (KSHV), the etiological agent of multiple human malignancies, including Kaposi’s sarcoma (KS), remains a significant concern, particularly in AIDS patients. A CRISPR-Cas9 screening of matched primary rat mesenchymal stem cells (MM) and KSHV-transformed MM cells (KMM) identified key splicing factors involved in KSHV-induced cellular transformation. To elucidate the mechanisms by which KSHV-driven splicing reprogramming mediates cellular transformation, we performed transcriptomic sequencing, identifying 131 differential alternative splicing transcripts, with exon skipping as the predominant event. Notably, these transcripts were enriched in vascular permeability, multiple metabolic pathways and ERK1/2 signaling cascades, which play key roles in KSHV-induced oncogenesis. Further analyses of cells infected with KSHV mutants lacking latent genes including vFLIP, vCyclin and viral miRNAs, as well as cells overexpressing LANA, revealed their involvement in alternative splicing regulation. Among the identified splicing factors, FAM50A, a component of the spliceosome complex C, was found to be crucial for KSHV-mediated transformation. FAM50A knockout resulted in distinct splicing profiles in both MM and KMM cells, and significantly inhibited KSHV-driven proliferation, cellular transformation and tumorigenesis. Mechanistically, FAM50A knockout altered SHP2 splicing, promoting an isoform with enhanced enzymatic activity that led to reduced STAT3 Y705 phosphorylation in KMM cells. These findings reveal a novel paradigm in which KSHV hijacks host splicing machinery, specifically FAM50A-mediated SHP2 splicing, to sustain STAT3 activation and drive oncogenic transformation.

**Importance:** Kaposi’s sarcoma-associated herpesvirus (KSHV) causes cancers such as Kaposi’s sarcoma, particularly in AIDS patients. This study uncovers how KSHV hijacks a fundamental cellular process called RNA splicing to promote cancer development. We identified key splicing events that alter critical pathways involved in vascular permeability, metabolism, and oncogenic signaling, particularly ERK1/2 and STAT3. A specific protein, FAM50A, was found to be essential for KSHV-driven cancerous transformation. Removing FAM50A disrupted splicing, weakening cancer-promoting signals. These findings provide new insights into how viruses manipulate host cells to drive cancer and highlight RNA splicing as a potential target for future therapies.

## Introduction

Alternative splicing is a fundamental post-transcriptional mechanism that regulates gene expression in eukaryotic cells (1). It relies on the precise recognition of splice sites and selective intron removal, orchestrated by the spliceosome, a dynamic ribonucleoprotein (RNP) complex consisting of five small nuclear RNAs (snRNAs; U1, U2, U4, U5, and U6) and numerous associated proteins (2). Through alternative splicing, a single precursor mRNA (pre-mRNA) can be differentially spliced to generate multiple mRNA isoforms, significantly expanding the proteomic diversity of eukaryotic organisms (2). This process is essential for the functional complexity of the mammalian proteome, allowing diverse biological functions to emerge from a relatively limited number of genes (3).

Approximately 90% of human pre-mRNAs undergo alternative splicing, producing mRNA isoforms that exhibit cell type-, tissue-, and developmental stage-specific expression patterns (4). However, dysregulation of alternative splicing has been strongly implicated in human diseases, particularly cancer (5). Comprehensive transcriptomic analyses have revealed that nearly all cancer tissues exhibit abnormal alternative splicing profiles compared to their normal counterparts (6–8). Increasing evidence suggests that tumor-specific splicing variants contribute to key oncogenic processes, including proliferation, invasion, metastasis, apoptosis evasion, drug resistance, and metabolic reprogramming (6–8). Given the profound impact of alternative splicing on gene regulation, understanding its molecular mechanisms and biological significance in both normal and cancerous cells is critical for advancing our knowledge of cell biology and cancer pathogenesis.

Kaposi’s sarcoma-associated herpesvirus (KSHV) is the etiological agent of multiple human malignancies, including Kaposi’s sarcoma (KS), primary effusion lymphoma (PEL), multicentric Castleman’s disease (MCD), and KSHV-associated inflammatory cytokine syndrome (KICS) (9–12). KSHV has a complex life cycle consisting of two distinct transcriptional programs: latency and the lytic phase. During latency, a restricted set of viral genes is expressed, primarily to maintain viral persistence and evade the host immune response (13). In contrast, the lytic phase involves the sequential expression of numerous viral genes, leading to viral replication and the production of infectious virions (14).

Previous studies have identified alternative splicing in multiple KSHV genes and highlighted the involvement of various viral and host factors in regulating these splicing events (15). However, whether KSHV actively reprograms the host cell alternative splicing landscape during infection and cellular transformation remains unclear.

To address this question, we previously established a KSHV-induced cellular transformation model using primary rat embryonic metanephric mesenchymal precursor (MM) cells, which can be efficiently infected and transformed by KSHV (16). Compared to untransformed MM cells, KSHV-transformed MM (KMM) cells exhibit characteristics of oncogenic transformation, including immortalization, enhanced proliferation, loss of contact inhibition, and tumorigenic potential in vivo (16). This unique system has been instrumental in identifying both viral and host factors that drive KSHV-mediated oncogenesis.

To uncover the cellular mechanisms underlying KSHV-driven cellular transformation, we previously performed a genome-wide CRISPR-Cas9 screening in matched MM and KMM cells (17). This analysis identified a set of genes that were essential for the survival of KMM but not MM cells, highlighting key cellular factors involved in KSHV-mediated oncogenesis. Given the limited studies on splicing regulation during KSHV infection and the lack of research on the mechanisms governing alternative splicing regulation of cellular genes and splicing factors (18), we sought to further characterize the essential cellular genes involved in KSHV-induced cellular transformation identified in our CRISPR-Cas9 screen.

The family with sequence similarity (FAM) gene group comprises multiple genes sharing conserved sequences and playing critical roles in various diseases (19, 20). Among them, the FAM50 gene family consists of two members: FAM50A and FAM50B (19, 20). FAM50A encodes a nuclear DNA-binding transcription factor involved in mRNA processing and has been identified as a splicing factor that interacts with spliceosome U5 and C-complex proteins (21). Mutations in FAM50A are linked to Armfield XLID syndrome, a spliceosomopathy characterized by defects in RNA splicing. Emerging evidence suggests that FAM50A functions as a proto-oncogene, contributing to the progression of multiple cancers. Elevated FAM50A expression correlates with poor prognosis and negative response to immunotherapy across several cancer types (22–24). However, despite its clinical significance, the precise role of FAM50A in oncogenesis and cancer progression remains largely unexplored.

Here, we map KSHV-driven alternative splicing events (ASEs) and the resulting alternatively spliced transcripts during KSHV-induced cellular transformation. We identify viral latent genes involved in this process and further investigate the role of FAM50A, an essential factor in KSHV-mediated cellular transformation, in regulating alternative pre-mRNA splicing. Specifically, we demonstrate that FAM50A modulates the alternative splicing of SHP2, generating distinct short and long isoforms that promote KSHV-induced oncogenesis. These findings highlight a critical function of FAM50A in KSHV-driven alternative splicing regulation, offering new insights into the molecular mechanisms underlying KSHV-associated malignancies.

## Results

### Splicing factors play an essential role in KSHV-induced cellular transformation

Alternative splicing is associated with the oncogenesis across various malignancies (25). To investigate alternative splicing regulation in KSHV-induced cellular transformation, we performed bulk RNA sequencing (RNA-seq) and identified 22 differentially expressed splicing factors in KSHV-transformed cells. Among them, six genes (JUP, UBL5, FAM50A, SAP30BP, EIF4A3 and GPATCH1) were upregulated, while 16 genes (PRPF18, CIRBP, RBMX, TXNL4A, KIN, LSM6, PABPC1, SNRPG, USP39, SNRNP27, BAG2, ISY1, SNRPB2, LSM3, DNAJC6 and KHDRBS3) were downregulated (Fig. 1A), suggesting KSHV might target these splicing factors.

**FIG 1.**
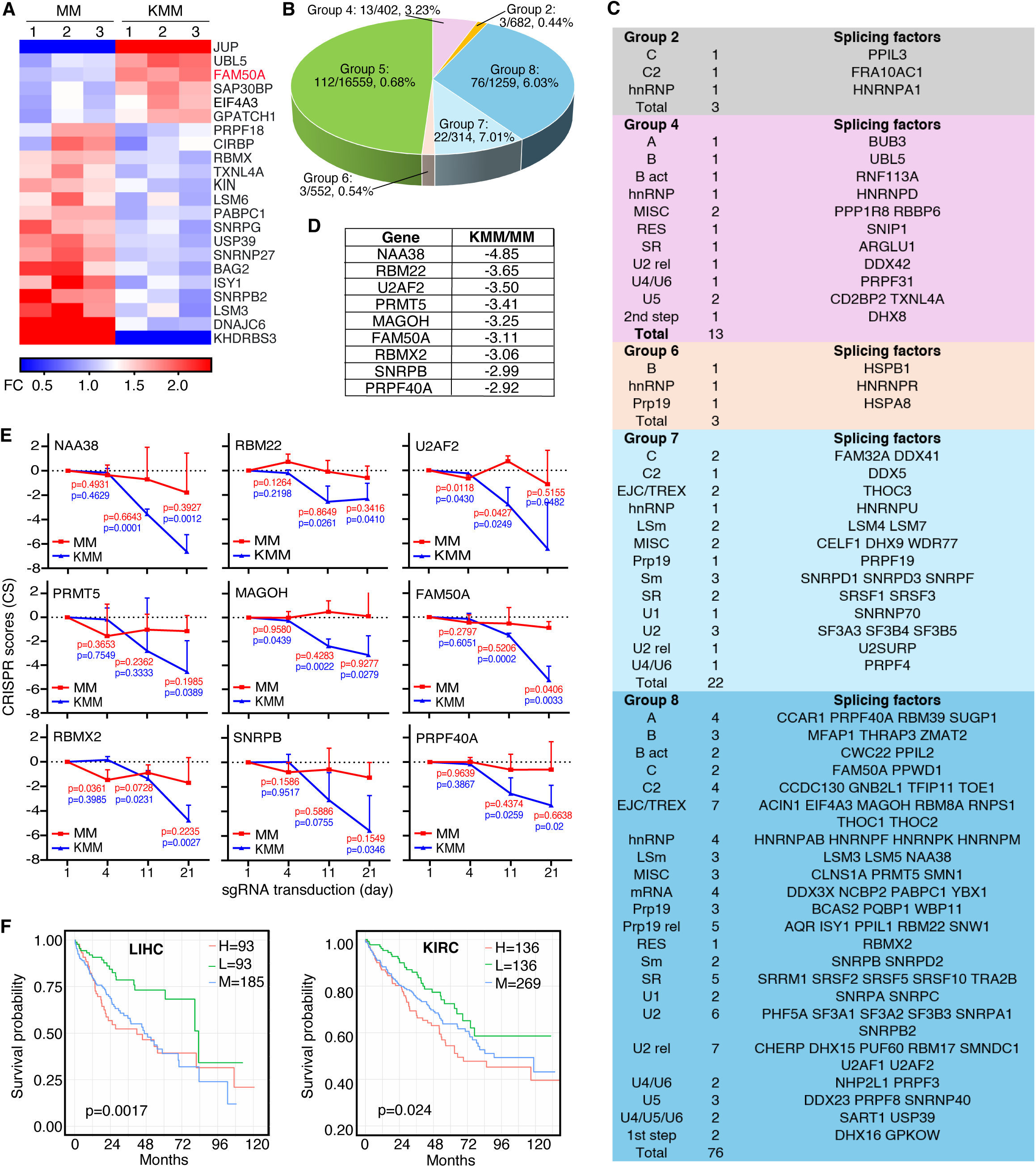
Splicing factors are essential for KSHV-induced cellular transformation. (A) Heatmaps showing the dysregulation of splicing factors in MM and KMM cells. (B) Distribution of splicing factors in different functional groups in MM and KMM cells identified by CRISPR-Cas9 screening (17). (C) Functional classification of splicing factors (26) identified as essential factors by CRISPR-Cas9 screening in MM and KMM cells. SR, serine-/arginine-rich protein; hnRNP, heterogeneous nuclear ribonucleoprotein; U2 rel, U2 related; Sm, small RNA-binding protein; LSm, like Sm; RES, RES complex protein; Bact, Bact complex protein; MISC, miscellaneous. (D) Top nine splicing factors essential for the proliferation of KMM cells with the largest differences in CRISPR scores between MM and KMM cells, identified in Group 8 in the screening. CRISPR scores represent the average [log_2_(final sgRNA abundance/initial sgRNA abundance)] of three sgRNAs. (E) CRISPR scores of the top nine splicing factors measured at day 4, 11, and 21 in MM (red) and KMM (blue) cells. *P*-values were obtained by comparing with day 1 for each cell type. (F) Survival analysis of FAM50A expression in liver hepatocellular carcinoma (LIHC) and kidney renal clear cell carcinoma (KIRC). H, M, and L represent high, medium, and low expression groups, respectively.

To further assess the functional significance of these splicing factors, we leveraged results from a CRISPR-Cas9 screening in KSHV-transformed (KMM) and untransformed (MM) cells (17). This screen identified over 200 splicing-related genes, with approximately 50% being essential for KMM cell proliferation and survival, but not for MM cells (Fig. 1B) (26). Notably, these essential genes were highly enriched in Groups 7 and 8 (17), which are essential for KMM cell proliferation and survival, where splicing factors were nearly 10-fold overrepresented compared to other groups. Importantly, the identified splicing factors spanned all major spliceosome complexes and splicing factor families (Fig. 1C).

The top nine splicing factors in Group 8, ranked by CRISPR scores, were NAA38, RBM22, U2AF2, PRMT5, MAGOH, FAM50A, RBMX2, SNRPB, and PRPF40A (Fig. 1D). These factors all belong to core spliceosomal complexes and play key roles in pre-mRNA splicing (Fig. 1C). Notably, knockout of any of these nine genes significantly inhibited KMM cell proliferation and survival, while exerting minimal effects on MM cells (Fig. 1E). Among them, FAM50A was the only splicing factor that was upregulated in KMM cells relative to MM cells (Fig. 1A), highlighting its unique role in KSHV-driven cellular transformation.

Given the essential role of these nine splicing factors in KSHV-induced cellular transformation, we assessed their clinical relevance by analyzing their prognostic significance in cancer using TCGA survival data. High expression of any of these factors was correlated with poor survival in multiple cancer types (Fig. 1F, Fig. S1, Table S1), including FAM50A in adenoid cystic carcinoma (ACC), liver hepatocellular carcinoma (LIHC), esophageal cancer (ESCA), kidney renal clear cell carcinoma (KIRC), sarcoma (SARC), kidney chromophobe carcinoma (KICH), acute myeloid leukemia (LAML), mesothelioma (MESO) and uveal melanoma (UVM) (Fig. S1A); NAA38 in colon adenocarcinoma (COAD), LIHC and MESO (Fig. S1B); RBM22 in KICH, LIHC, kidney renal papillary cell carcinoma (KIRP) and SARC (Fig. S1C); U2AF2 in ACC, brain lower grade glioma (LGG), LAML, MESO, LIHC, SARC and UVM (Fig. S1D); PRMT5 in bladder urothelial carcinoma (BLCA), LIHC, head and neck squamous cell carcinoma (HNSC) and thyroid carcinoma (THCA) (Fig. S1E); MAGOH in ACC, LIHC, LGG, KIRP, MESO and SARC (Fig. S1F); RBMX2 in ESCA, LIHC, HNSC, KIRC, KIRP and uterine corpus endometrial carcinoma (UCEC) (Fig. S1G); SNRPB in LGG, LIHC, COAD, KIRC, KIRP, LAML, lung squamous cell carcinoma (LUSC), MESO, SARC and UVM (Fig. S1H); and PRPF40A in ACC, LIHC, LGG, KICH, KIRP, lung adenocarcinoma (LUAD), MESO, pancreatic adenocarcinoma (PAAD) and UCEC (Fig. S1I).

These findings demonstrate that KSHV-induced cellular transformation profoundly reshapes alternative splicing regulation, highlighting the critical role of splicing factors in oncogenesis. Among the identified factors, FAM50A emerges as a key mediator of KSHV-driven splicing reprogramming, potentially contributing to the pathogenesis of KSHV-associated malignancies.

### KSHV reprograms alternative pre-mRNA splicing through viral latent genes and miRNAs

To investigate how KSHV infection regulates host cell alternative splicing, we performed splicing analysis using SUPPA2 (27) and identified 131 differential spliced transcripts between MM and KMM cells (Fig. 2A). Among these, cassette exon (SE) events accounted for 68%, while alternative first exon (AFE) events represented 19% (Fig. 2B). Gene ontology (GO) analysis of biological processes (BP) revealed that the affected transcripts were highly enriched in pathways implicated in KSHV infection and cellular transformation, including the regulation of vascular permeability (16, 28, 29), multiple metabolic pathways (30–32), and ERK1/2 signaling cascades (33–40) (Fig. 2C).

**FIG 2.**
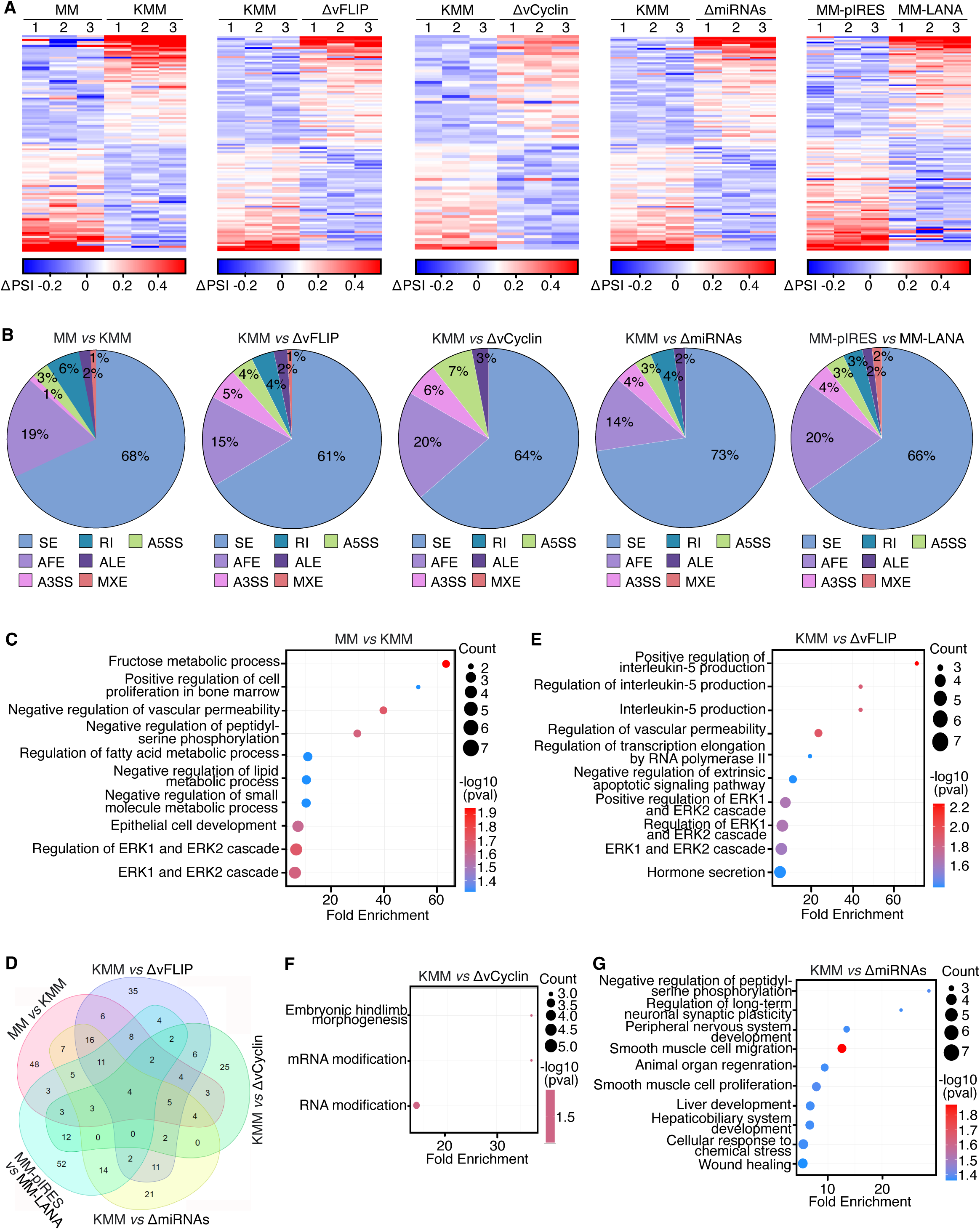
KSHV latent genes and miRNAs contribute to splicing alterations in KSHV-transformed cells. (A) Differential spliced transcripts between MM vs. KMM, KMM vs. ΔvFLIP, KMM vs. ΔvCyclin, KMM vs. ΔmiRNAs, and MM-pIRES vs. MM-LANA, measured as ΔPSI values from three independent biological replicates. (B) Pie charts depicting ASE types classified by SUPPA2 between MM vs. KMM, KMM vs. ΔvFLIP, KMM vs. ΔvCyclin, KMM vs. ΔmiRNAs, and MM-pIRES vs. MM-LANA. Chart size represents the number of differential spliced transcripts detected. (C) GO enrichment analysis of biological processes (BPs) affected by differential spliced transcripts between MM vs. KMM. (D) Venn diagram summarizing overlapping differential spliced transcripts across all comparisons. (E-G) GO enrichment analysis of differential spliced transcripts between KMM vs. ΔvFLIP (E), KMM vs. ΔvCyclin (F), and KMM vs. ΔmiRNAs (G).

During latent infection, KSHV expresses only a limited set of viral genes, including LANA, vFLIP, vCyclin, and a cluster of viral miRNAs (14, 41). To determine whether these latent genes contribute to the regulation of host alternative splicing, we performed splicing analysis in MM cells infected with KSHV mutants lacking vFLIP (ΔvFLIP), vCyclin (ΔvCyclin), or a cluster of pre-miRNAs (ΔmiRNAs) (Fig. 2A-B). Since LANA is essential for latent infection (42, 43), we generated MM cells expressing LANA (MM-LANA) and compared their splicing profiles with vector control (MM-pIRES) cells. Across all conditions, SE events were the most prevalent ASEs, constituting 61-73% of the total (Fig. 2B).

A Venn diagram analysis revealed that vFLIP and KSHV miRNAs had a more substantial impact on KSHV-induced alternative splicing reprogramming than vCyclin and LANA (Fig. 2D). Specifically, 50.4% (63/125) of differential spliced transcripts in ΔvFLIP cells and 42.4% (55/105) in ΔmiRNAs cells overlapped with those in KMM cells. In contrast, only 37.3% (28/75) of differential spliced transcripts in ΔvCyclin cells and 31.2% (39/125) in LANA-overexpressing cells overlapped with KMM cell transcripts (Fig. 2D).

GO analysis further confirmed that transcripts differentially spliced in ΔvFLIP cells were enriched in pathways related to vascular permeability, ERK1/2 cascades, transcription elongation by RNA polymerase II, and interleukin-5 (IL-5) signaling (Fig. 2E). Similarly, transcripts differentially spliced in ΔmiRNAs cells were enriched in pathways associated with cell proliferation, migration, wound healing, and stress response (Fig. 2G). In contrast, few (3) enriched pathways were found for differential spliced transcripts in ΔvCyclin cells, of which 2 are related to RNA modification (Fig. 2F), while no significantly enriched pathways were identified in LANA-overexpressing cells (data no shown).

Together these findings suggest that KSHV reprograms host alternative splicing primarily through vFLIP and viral miRNAs, highlighting their pivotal roles in KSHV-induced oncogenic transformation (32, 44).

### FAM50A is essential for KSHV-induced cellular transformation and tumorigenesis

RNA-seq analysis revealed that FAM50A is significantly upregulated in KMM cells compared to MM cells (Fig. 1A and 3A), a finding further validated by Western blot analysis (Fig. 3B). Similarly, elevated FAM50A expression was observed in KSHV-infected PEL cell lines (BC3 and BCP1) and KSHV-infected BJAB cells (BJAB-KSHV) compared to uninfected BJAB cells (Fig. 3C).

**FIG 3.**
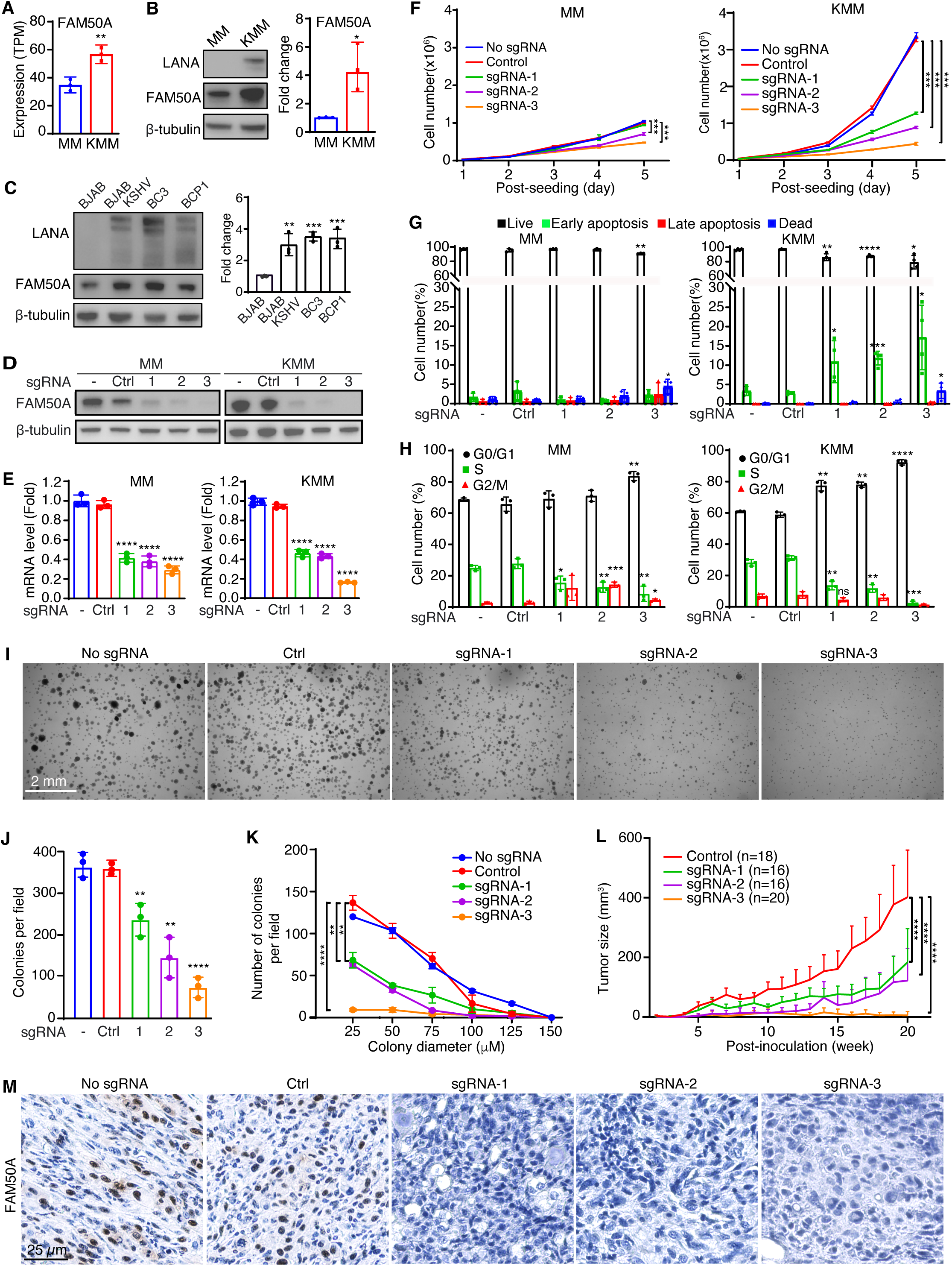
FAM50A is essential for KSHV-induced cellular transformation. (A) Relative mRNA expression of FAM50A in MM and KMM cells, measured by RNA-seq (TPM, transcripts per million). (B) Western blot analysis of FAM50A expression in MM and KMM cells and quantifications from three independent experiments. (C) Immunoblotting analysis and quantifications of FAM50A expression in PEL, BJAB-KSHV and BJAB cells. (D-E) Validation of FAM50A knockout by Western blotting (D) and RT-qPCR (E). (F-H) Effects of FAM50A knockout on cell proliferation (F), apoptosis (G), and cell cycle progression (H) in MM and KMM cells. (I-K) FAM50A knockout reduced colony formation efficiency in KMM cells grown on soft agar (I) quantified in colony numbers (J) and size distribution (K). (L) FAM50A knockout suppresses KMM tumor progression in nude mice. *P*-values were obtained by comparing tumor volumes at the 20-week endpoint between the indicated groups. (M) IHC staining of FAM50A in KS-like tumors from nude mice. Data are presented as mean ± 95% CI and *P*-values (\**P* < 0.05; \*\**P* < 0.01, \*\*\**P* < 0.001, \*\*\*\**P* < 0.0001) were determined using Student’s *t*-test.

To determine the functional role of FAM50A in KSHV-induced cellular transformation, we generated FAM50A knockout cell lines in both MM and KMM cells (Figs. 3D-E). FAM50A knockout significantly reduced the proliferation of KMM cells but had only a minor effect on MM cells (Fig. 3F). Apoptosis analysis revealed that FAM50A knockout triggered early apoptosis in KMM cells while having minimal impact on MM cells (Fig. 3G). Additionally, cell cycle analysis demonstrated that FAM50A knockout induced G0/G1 cell cycle arrest in both MM and KMM cells, with a more pronounced effect in KMM cells (Fig. 3H). These findings suggest that FAM50A plays a key role in promoting KSHV-induced cell proliferation and survival.

To further assess the role of FAM50A in KSHV-driven cellular transformation, we conducted soft agar colony formation assay. FAM50A knockout significantly impaired the ability of KMM cells to form colonies, reducing both colony number and size (Fig. 3I-K). To evaluate its role in tumorigenesis, we examined tumor growth in nude mice using FAM50A knockout cells. Loss of FAM50A markedly suppressed tumor progression in KMM-derived xenografts (Figs. 3L and S2). Immunohistochemical analysis confirmed strong FAM50A expression in tumors derived from KMM-Cas9 cells (No sgRNA) and KMM cells with scrambled sgRNAs (Control), whereas tumors from FAM50A knockout cells showed no detectable expression (Fig. 3M). Together, these results demonstrate that FAM50A plays a critical role in KSHV-induced cellular transformation and tumorigenesis.

### FAM50A regulates alternative pre-mRNA splicing in MM and KMM cells

Given that FAM50A functions as an essential splicing factor in KMM cells, we investigated the alternative spliced transcripts regulated by FAM50A in MM and KMM cells. RNA-seq analysis of FAM50A knockout MM and KMM cells identified 34 differential spliced transcripts in MM cells and 28 in KMM cells (Fig. 4A-B and S3). GO analysis revealed that both MM and KMM cells share common alternative spliced transcripts upon FAM50A knockout, including those related to fructose metabolism and carbohydrate phosphorylation (Fig. 4C-D). However, alternative spliced transcripts in FAM50A knockout KMM cells were also enriched in pathways crucial for KSHV-induced cellular transformation, such as apoptosis, oxidative stress, and platelet-derived growth factor (PDGF) signaling (Fig. 4D). While 16 of the 34 differential spliced transcripts in MM FAM50A knockout cells and 12 of the 28 in KMM FAM50A knockout cells overlapped with those in KMM cells, only four were common across both cell types (Fig. 4E). This suggests that FAM50A regulates distinct ASEs in MM and KMM cells, potentially contributing to differences in biological functions.

**FIG 4.**
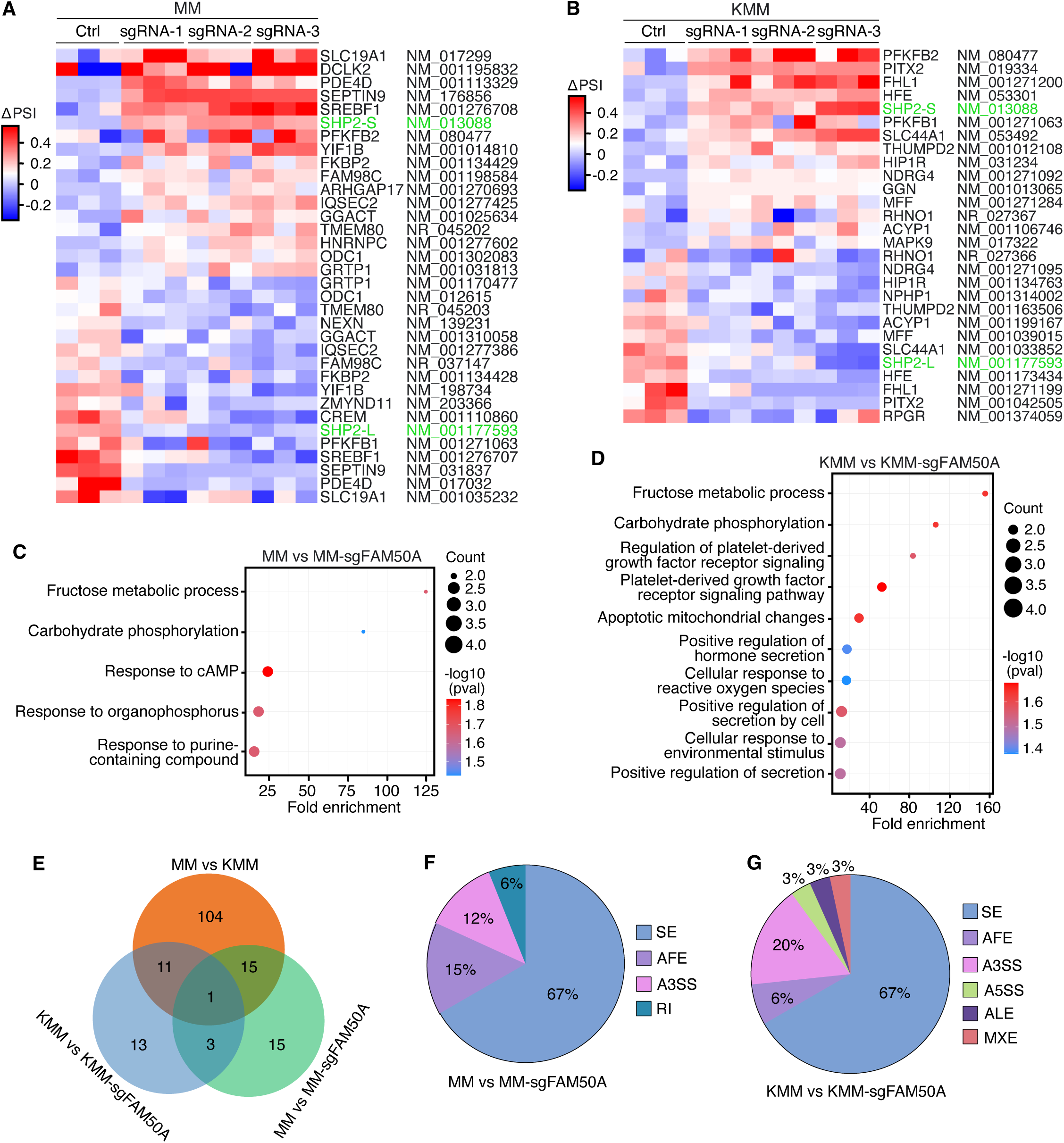
FAM50A knockout alters alternative RNA splicing in primary and KSHV-transformed cells. (A-B) Heatmaps showing ΔPSI values for differential spliced transcripts between MM vs. MM-sgFAM50A (A) and KMM vs. KMM-sgFAM50A (B) cells, based on three independent biological replicates. (C-D) GO enrichment analysis of differential spliced transcripts in MM-sgFAM50A (C) and KMM-sgFAM50A (D) cells. (E) Venn diagram depicting the overlaps of differential spliced transcripts between MM vs. KMM, MM vs. MM-sgFAM50A and KMM vs. KMM-sgFAM50A cells. (F and G) Pie charts showing the distribution of ASE types in MM vs. MM-sgFAM50A (F) and KMM vs. KMM-sgFAM50A (G) groups, classified by SUPPA2.

Among the various types of alternative splicing, SE events were the most predominant, accounting for 67% in both FAM50A knockout MM and KMM cells (Figs. 4F-G). Notably, alternative 3’ splice site selection (A3SS) accounted for 20% of differential spliced transcripts in KMM FAM50A knockout cells, but only 12% in MM FAM50A knockout cells.

Collectively, these findings indicate that FAM50A is a key regulator of alternative pre-mRNA splicing in both MM and KMM cells, with distinct functional consequences in KSHV-transformed cells.

### FAM50A enhances STAT3 activation and cell proliferation by regulating SHP2 alternative splicing

To determine which alternative spliced transcripts induced by FAM50A knockout contribute to the inhibition of KMM cell proliferation, we focused on the most differential spliced transcripts following FAM50A knockout in MM and KMM cells (Fig. 4A-B). Among all identified alternative spliced transcripts, SHP2 transcripts were the most significantly affected in both MM and KMM cells following FAM50A knockout. SHP2 encodes two isoforms: a short isoform (SHP2-S) and a long isoform (SHP2-L). While FAM50A knockout did not alter the total SHP2 expression level, it upregulated SHP2-S while downregulating SHP2-L (Fig. 5A and S4A). These results were further validated by semi-quantitative RT-PCR (Fig. 5B-C and S4B).

**FIG 5.**
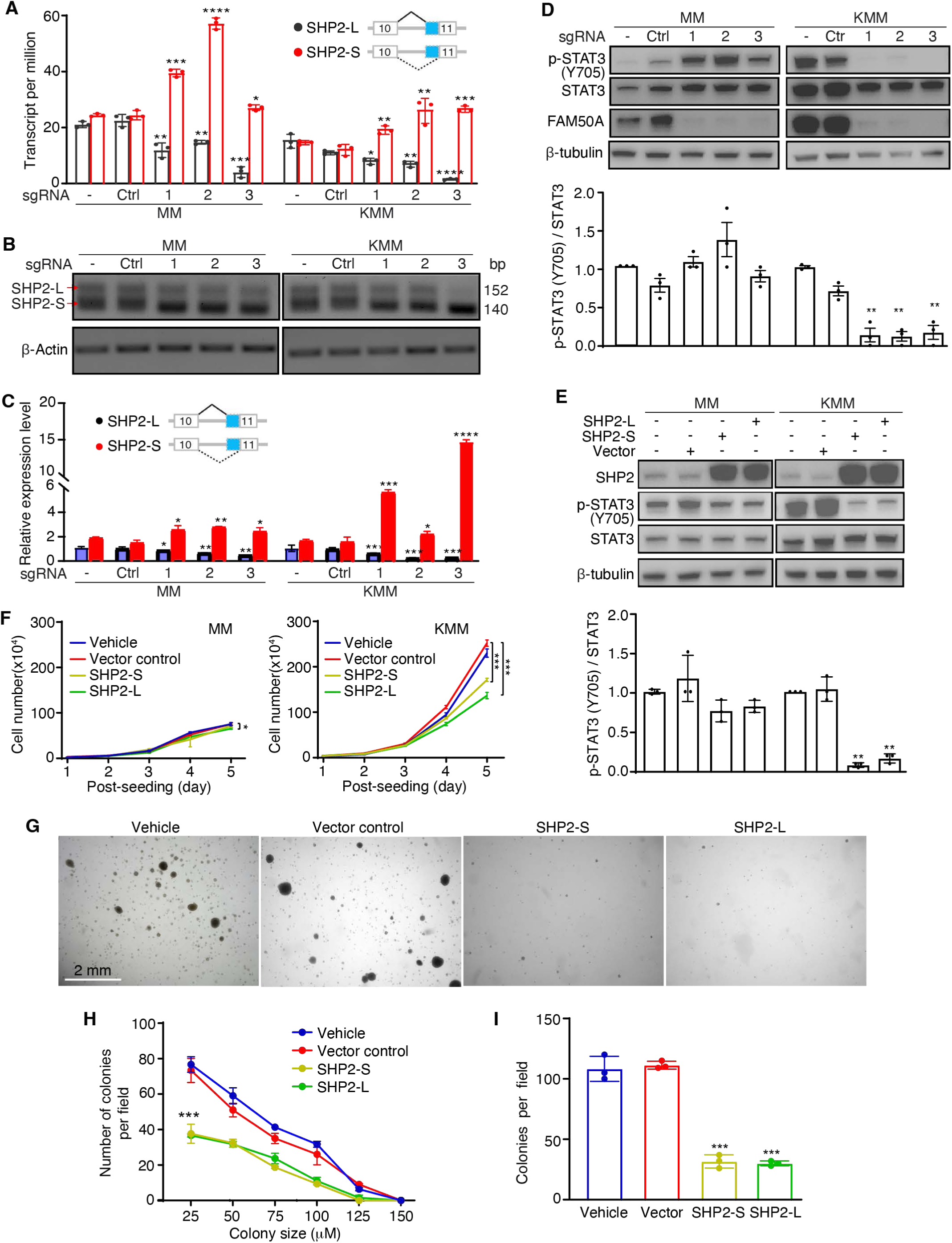
FAM50A regulates alternative splicing of SHP2 isoforms to activate STAT3 pathway in KSHV-transformed cells. (A) RNA-seq analysis reveals that SHP2-long (SHP2-L) isoform expression decreases while SHP2-short (SHP2-S) isoform expression increases following FAM50A knockout in MM and KMM cells. (B-C) Validation of SHP2 isoform switching by RT-PCR and gel electrophoresis (B) and quantification of the relative SHP2-S/L intensity (C). (D) Western blot analysis of STAT3 signaling in MM and KMM cells after FAM50A knockout with lower panel showing the quantifications of the relative p-STAT3 (Y705) levels. (E) Immunoblotting of STAT3 signaling in SHP2-S overexpressing (SHP2-S-OE) and SHP2-L overexpressing (SHP2-L-OE) MM and KMM cells, with lower panel showing quantifications of the relative p-STAT3 (Y705) levels. (F) Growth curves of SHP2-S-OE and SHP2-L-OE MM and KMM cells. (G-I) Soft agar assay demonstrating the impact of overexpressing SHP2-S and SHP2-L on colony formation of KMM cells (G) with quantifications of colony numbers (H) and size distribution (I). Data are presented as mean ± 95% CI and *P*-values (\**P* < 0.05; \*\**P* < 0.01, \*\*\**P* < 0.001, \*\*\*\**P* < 0.0001) were determined using Student’s *t*-test.

Given that SHP2-S exhibits 10-fold higher phosphatase catalytic activity than SHP2-L (45) and that FAM50A knockout leads to an increase in SHP2-S, we hypothesized that SHP2 phosphatase activity could be elevated following FAM50A knockout. To test this, we examined the phosphorylation of STAT3-Y705, a known SHP2 downstream target (46). Indeed, FAM50A knockout significantly inhibited STAT3-Y705 phosphorylation in KMM cells (Fig. 5D). Unexpectedly, STAT3-Y705 phosphorylation was increased in MM cells following FAM50A knockout, suggesting that SHP2 regulation of STAT3 activation is cell type-dependent.

To further investigate this, we overexpressed SHP2-S or SHP2-L in MM and KMM cells. Both SHP2-S and SHP2-L inhibited STAT3 activation in MM and KMM cells, but the inhibitory effect was more pronounced in KMM cells (Fig. 5E). Consistently, overexpression of SHP2-S or SHP2-L inhibited KMM cell proliferation, whereas MM cell proliferation was largely unaffected (Fig. 5F). Moreover, SHP2-S or SHP2-L overexpression suppressed colony formation of KMM cells, leading to a reduction in both colony number and size (Fig. 5G-I).

To further confirm the role of SHP2 in mediating FAM50A-dependent STAT3 regulation, we performed siRNA-mediated SHP2 knockdown in FAM50A knockout KMM cells. SHP2 depletion restored STAT3-Y705 phosphorylation (Fig. 6A-B), reinforcing the notion that SHP2 mediates STAT3 inhibition in FAM50A knockout KMM cells. However, SHP2 knockdown failed to rescue the inhibition of cell proliferation caused by FAM50A knockout (Fig. 6C). In line with this observation, SHP2 knockdown did not restore colony formation in soft agar following FAM50A depletion (Fig. 6D). These results suggest that STAT3 activation and cell proliferation are regulated by additional factors besides SHP2, implying that SHP2 is not the sole downstream target of FAM50A.

**FIG 6.**
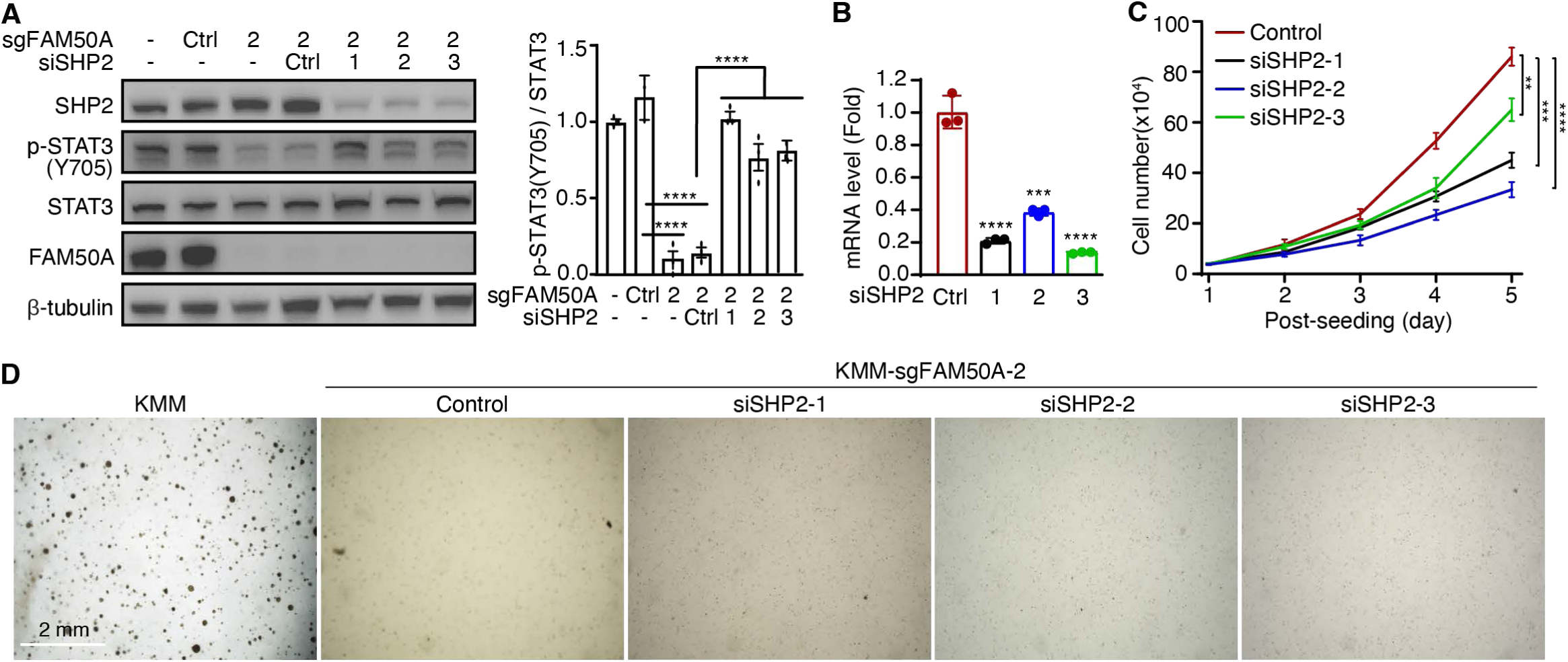
SHP2 knockdown partially recues p-STAT3 activation in FAM50A knockout KMM cells. (A) Immunoblot analysis of STAT3 activation in FAM50A knockout KMM cells following SHP2 knockdown with quantifications shown in the right panel. (B) Confirmation of SHP2 knockdown efficiency in FAM50A knockout KMM cells by RT-qPCR. (C) Growth curves of FAM50A knockout KMM cells with or without SHP2 knockdown. (D) Soft agar colony formation assay of FAM50A knockout KMM cells after SHP2 knockdown. Data are presented as mean ± 95% CI and *P*-values (\**P* < 0.05; \*\**P* < 0.01, \*\*\**P* < 0.001, \*\*\*\**P* < 0.0001) were determined using Student’s *t*-test.

Together, our findings indicate that FAM50A plays a critical role in KSHV-induced cell proliferation and cellular transformation by regulating SHP2 alternative splicing and STAT3 activation.

## Discussion

Understanding how cancer cells reprogram splicing to facilitate oncogenesis is fundamental to cancer biology. In this study, we uncovered that oncogenic KSHV manipulates multiple splicing factors to promote the proliferation and survival of KSHV-transformed cells. Our findings demonstrate that KSHV systemically reprograms alternative splicing in host cells during cellular transformation and that KSHV latent genes, particularly vFLIP and KSHV miRNAs, play a crucial role in this process. While extensive research has focused on how KSHV latent genes mediate viral latent replication and transcriptional regulation, their roles in alternative splicing have remained largely unexplored (14). Here, we provide compelling evidence that KSHV infection and latent genes contribute to splicing regulation, a novel aspect of KSHV-driven oncogenesis. Further studies are warranted to determine whether these findings extend to other KSHV-latently infected cell types.

Although the precise mechanisms by which KSHV infection and latent genes regulate alternative splicing remain to be fully elucidated, our results indicate that KSHV alters the expression of multiple splicing factors in transformed cells (Fig. 1A). Beyond transcriptional control, KSHV latent proteins may directly interact with RNA-binding proteins or act as splicing regulators by associating with pre-mRNA or the spliceosome, analogous to the function of the KSHV lytic protein ORF57 (47). Notably, the alternative spliced transcripts modulated by KSHV latent genes only partially overlap with those driven by KSHV-induced cellular transformation (Fig. 2D), suggesting that additional factors contribute to splicing dysregulation. While some alternative spliced transcripts are shared, each KSHV latent gene appears to regulate a distinct subset, underscoring the complexity and specificity of host splicing alterations during KSHV-induced cellular transformation.

Aberrant splicing is a well-documented hallmark of cancer, with specific splicing factors implicated in tumorigenesis (25). Alternative splicing contributes to transcriptomic diversity, affecting key cellular processes such as proliferation, apoptosis, and immune evasion (5). However, little is known about how oncogenic viruses alter alternative splicing and splicing factors in virus-induced cancers. To address this gap, we analyzed results of our previous genome-wide CRISPR-Cas9 screen to systematically identify splicing factors essential for KSHV-induced cellular transformation (Fig. 1C-D) (17). Among these factors, FAM50A emerged as a critical regulator, exhibiting significant upregulation in KSHV-transformed cells. Unlike well-characterized spliceosomal components, FAM50A remains largely unexplored in both cancer and viral infection. Our study has shown that FAM50A is not only upregulated in KSHV-transformed cells but also actively participates in alternative splicing regulation. These results align with prior reports describing the oncogenic roles of other spliceosomal proteins (48, 49).

Importantly, we identified FAM50A as an essential factor in KSHV-induced cellular transformation, functioning in part by modulating KSHV-driven splicing reprogramming (Fig. 3). FAM50A knockout selectively alters SHP2 isoform expression, shifting it toward a catalytically active short isoform in KSHV-transformed cells (Fig. 5A-C) (45). This isoform switch results in decreased phosphorylation of STAT3 at Y705 (Fig. 5D), a key driver of KSHV-induced cellular transformation (30, 50, 51). Consequently, FAM50A deficiency suppresses KSHV-transformed cell proliferation and anchorage-independent growth in soft agar (Fig. 5F-I). These findings implicate FAM50A as a virus-specific regulator of SHP2 catalytic activity, revealing a novel mechanism by which KSHV exploits the host splicing machinery to promote oncogenesis.

The identification of FAM50A as an essential factor for KSHV-induced transformation suggests its potential as a therapeutic target. Given that FAM50A is associated with poor prognosis in multiple cancers (Fig. 1F and S1A) (22), its inhibition could have broad therapeutic implications (52). Our findings indicate that FAM50A’s function differs between primary and KSHV-transformed cells (Fig. 5D-F), raising the possibility of selectively targeting FAM50A in virus-associated malignancies. Future research should explore potential strategies for FAM50A inhibition, including small-molecule inhibitors or RNA-based therapeutics, and evaluate possible off-target effects in non-cancerous cells.

While our study provides novel insights into FAM50A’s role in KSHV-induced cellular transformation, several limitations should be acknowledged. First, our findings are based primarily on in vitro cell culture and immunocompromised mouse models; further validation in KSHV-induced human cancers is necessary. Second, although FAM50A has been proposed as a promising therapeutic target (22, 23), its clinical potential remains unexplored, necessitating further studies to assess the safety and efficacy of its inhibition. Additionally, an intriguing question remains as to whether KSHV latent proteins directly interact with FAM50A to regulate alternative splicing. Given that nuclear-localized KSHV latent proteins, such as LANA, may co-reside with spliceosomal components, they may directly interact with FAM50A and other splicing factors to mediate splicing regulation. Future studies should investigate these interactions and identify the essential ASEs driving KSHV-induced transformation.

In summary, our study demonstrates that splicing factors play fundamental roles in KSHV-induced cellular transformation. KSHV systematically reprograms host splicing, with its latent genes including vFLIP, vCyclin, LANA, and miRNAs closely involved in alternative splicing regulation. Furthermore, we identified FAM50A as a key splicing factor that promotes KSHV-induced cellular transformation by modulating alternative splicing and shifting SHP2 isoform expression, leading to decreased STAT3-Y705 phosphorylation. These findings reveal a novel mechanism by which KSHV manipulates the host splicing machinery to drive oncogenesis. Targeting FAM50A may offer new therapeutic opportunities for KSHV-associated malignancies and potentially other cancers dependent on aberrant splicing.

## Materials and Methods

### Antibodies

The antibodies used for immunoblotting included rabbit anti-FAM50A/XAP5 (Abcam, ab186410), rat anti-LANA (Abcam, ab4103), rabbit anti-SHP2 (Cell Signaling Technology, 3397), mouse anti-STAT3 (Cell Signaling Technology, 9139), rabbit anti-phospho-STAT3 (Tyr705) (Cell Signaling Technology, 9145), mouse anti-FLAG (Sigma-Aldrich, F1804), and mouse anti-β-tubulin (Sigma-Aldrich, 7B9). For immunohistochemistry (IHC), a rabbit anti-FAM50A antibody (Abcam, ab186410) was used.

### Cell culture

MM and 293T cells were maintained in Dulbecco’s Modified Eagle Medium (DMEM) (Genesee, 25-500) supplemented with 10% fetal bovine serum (FBS) (Sigma-Aldrich, F2442) and 1% penicillin/streptomycin (Gibco, 15140-122) in a 5% CO_2_ incubator at 37°C. KMM cells were cultured under the same conditions as MM cells but with the addition of 250 μg/ml hygromycin. MM and KMM cells with stable FAM50A knockout were maintained in their respective media. Cells with stable SHP2-S or SHP2-L expression were cultured in their respective media supplemented with 10 μg/ml Blasticidin (ThermoFisher Scientific, A1113903).

The PEL cell lines (BC3 and BCP-1), the Burkitt lymphoma cell line (BJAB), and the KSHV-infected BJAB cell line (BJAB-KSHV) were cultured in RPMI 1640 medium (Invitrogen, Carlsbad, CA) containing 20% FBS at 37°C under 5% CO_2_. Additionally, BJAB-KSHV cells were maintained in medium supplemented with 10 μg/ml Puromycin (Sigma-Aldrich, P8833).

All cell lines were cultured in drug-free medium for one week prior to experiments.

### Plasmids and transfection

All plasmids were constructed using the restriction enzyme digestion and ligation method. The pBSD-FLAG-SHP2-S and pBSD-FLAG-SHP2-L plasmids were generated by inserting PCR-amplified products into the pBSD vector using the EcoRI and XbaI restriction sites. The pBSD plasmid was obtained from Addgene (Plasmid #119863).

Plasmid transfection was carried out using Lipofectamine 2000 (Invitrogen, 11668019) following the manufacturer’s protocol. Cells were transfected using a ratio of 1 μg of plasmid DNA to 3 μl of Lipofectamine 2000 and cultured for three days before being used for the experiments.

### Generation of CRISPR/Cas9-mediated FAM50A gene knockout cell lines

FAM50A knockout KMM/MM cell lines were generated using the CRISPR/Cas9 (clustered regularly interspaced palindromic repeats/CRISPR-associated protein 9) system, following previously established protocols (17). Briefly, a single-guide RNA (sgRNA) sequence targeting the FAM50A locus (sgFAM50A sequence: 5’- TGGGCACCGGCGCACTGTTAAGG-3’) was designed on the basis of Cas-OFFinder (53) and Cas-Designer (54). The sgFAM50A plasmid was transduced into KMM/MM cells using lentivirus. The production of sgFAM50A lentivirus was performed as previously described (17).

Following transduction, cells were cultured for 48 hours before being subjected to stepwise serial dilution to isolate single-cell clones. Clones exhibiting puromycin resistance were selected and further analyzed by genomic DNA sequencing. Only clones displaying a single sequencing peak with a gap, indicative of successful gene knockout, were selected for subsequent experiments.

### Small interfering RNA-mediated SHP2 knockdown

Small interfering RNAs (siRNAs) targeting Rattus norvegicus SHP2 (SASI_Rn01_00105327, SASI_Rn01_00105328, SASI_Rn01_00105329) and the scramble control (SIC001) were purchased from Sigma-Aldrich. Transfection was performed using the Lipofectamine 2000 Kit (Invitrogen, 11668019) according to the manufacturer’s instructions. Cells were harvested 48 hours post-transfection to assess knockdown efficiency.

### RT-qPCR detection of gene expression

Total RNA was extracted from cultured cells using the TRIzol reagent (Sigma-Aldrich, T9424). First-strand cDNA synthesis was carried out with 50 to 100 ng of RNA per reaction using the Maxima H Minus First Strand cDNA Synthesis Kit (ThermoFisher Scientific, K1652). Quantitative real-time PCR (qPCR) was carried out using the SsoAdvanced Universal SYBR Green Supermix Kit (Bio-Rad, 172-5272) according to the manufacturer’s instructions. Gene expression levels were normalized to β-actin mRNA. The primers used for qPCR are listed in Table S2.

### Validation of ASEs by semiquantitative RT-PCR

Validation of alternative spliced transcripts was performed using semiquantitative RT-PCR. Total RNA was isolated, and first-strand cDNA was synthesized as described previously. PCR amplification was carried out using the Platinum™ PCR SuperMix High Fidelity (ThermoFisher Scientific, 12532016). Reaction products were separated on 2% agarose gels and visualized by ethidium bromide staining. The relative abundance of each splicing isoform was quantified using ImageJ software. Primers used for splicing assays are listed in Table S2.

### Cell proliferation assay

Cells were seeded into 12-well plates at a density of 30,000 cells per well. Three biological replicates were performed for each condition. At the indicated time points, cells were harvested and counted using a hemocytometer.

### Cell cycle and apoptosis assays

Cells were seeded into 6-well plates and cultured overnight. For cell cycle analysis, 5-bromo-2′-deoxyuridine (BrdU, 10 μM; Sigma-Aldrich, B5002) was added to the culture medium at 10 μM for 2 hours to label replicating DNA. Cells were then fixed with 70% ethanol, permeabilized with 2 M hydrochloric acid, and stained with an anti-BrdU monoclonal antibody (ThermoFisher Scientific, B35129). Cells were further stained with propidium iodide (PI), and flow cytometry was performed using a FACS Canto II system (BD Biosciences). Apoptotic cells were detected by flowcytometry using Fixable Viability Dye eFluor 660 (Invitrogen, 65-0864) and Annexin V Apoptosis Detection Kits (Invitrogen, 88-8103-74) according to the manufacturer’s instructions. Cells treated with 100 μM Menadione (Sigma-Aldrich, M5625) served as positive controls for apoptosis induction. All experiments were performed in three biological replicates, and data were analyzed using FlowJo software (BD Biosciences).

### Soft agar assay

The soft agar assay was performed as previously described (16). Briefly, 5 × 10^4^ cells were suspended in 1 mL of 0.3% top agar (Sigma-Aldrich, A5431) and plated onto a solidified 0.5% base agar layer in 6-well plates. The wells were then overlaid with culture medium to maintain cell viability. After 2 weeks of incubation, colonies were visualized using a microscope with a 2× objective lens, and colonies with a diameter greater than 50 μm were counted.

### Animal studies

All animal experiments were conducted under protocol 21079422, approved by the Institutional Animal Care and Use Committee (IACUC) at the University of Pittsburgh. Six-week-old male *Hsd:Athymic Nude-Foxn1^nu* mice (Envigo, Inotiv) were maintained under standardized pathogen-free conditions. FAM50A knockout KMM cells (sgFAM50A-1, 2 and 3), KMM-Cas9 cells, and KMM cells with scrambled sgRNAs (Control) were subcutaneously (IC) injected into both flanks of the mice at a density of 5 × 10^6^ cells in 0.1 ml of PBS per site using a 25-gauge needle. Tumor size was monitored twice weekly. The mice were euthanized at 20 weeks post-injection, and the tumors were excised, weighted and processed for immunohistochemistry.

### Immunohistochemistry staining

Formalin-fixed, paraffin-embedded (FFPE) tissue sections were rehydrated through a series of xylene and ethanol washes (100%, 95%, and 75%), followed by rinsing in water. Antigen retrieval was performed by pressure cooking the sections in citrate buffer at 110°C for 20 minutes. After washing to remove citrate buffer, endogenous peroxidase activity was quenched with 3% hydrogen peroxide in methanol for 30 minutes at room temperature. Sections were blocked with 5% BSA in 1× TBST for 1 hour at 37°C, followed by overnight incubation with the primary antibody (1:100 dilution in 2.5% BSA in 1× TBST) in a humidity-controlled chamber. The secondary antibody was applied at a 1:100 dilution in 2.5% BSA in 1× TBST and incubated for 60 minutes at 37°C. Slides were developed using the ImmPACT DAB Substrate Kit, Peroxidase (Vector, SK-4105), according to the manufacturer’s instructions, and counterstained with hematoxylin QS (Vector, H-3404-100). Finally, sections were dehydrated through graded ethanol (75%, 95%, and 100%) and xylene washes before being mounted with xylene-based mounting media (Epredia™ Shandon-Mount, 1900333).

### Analysis of splicing factors identified by CRISPR-Cas9 screening

Our prior CRISPR-Cas9 screening of MM and KMM cells identified essential genes for cell proliferation and survival (17). From this dataset, we mapped over 200 splicing-related factors (26) into nine functional groups based on the CRISPR-Cas9 screening results. The percentage of enrichment for each group was calculated, and splicing factors were further categorized based on their roles within the spliceosome.

### RNA-seq

Total RNA was extracted from cells using TRI Reagent (Sigma-Aldrich, T9424) following the manufacturer’s instructions. Approximately 10 μg of RNA from each sample was used for mRNA library preparation. The TruSeq Stranded mRNA Library Prep Kit (Illumina) was used to construct next-generation sequencing libraries according to the manufacturer’s protocol. Paired-end sequencing (150 base pairs) was performed on an Illumina HiSeq 4000 platform (Illumina).

### RNA-seq data analysis

RNA-seq data analysis was performed by first processing raw reads in fastq format to obtain clean reads, which were then aligned to the rat genome mRatBN7.2 using STAR (55). Transcripts were assembled, and raw gene counts were estimated using featureCounts (56) and presented in TPM (transcripts per million). Differentially expressed genes (DEGs) were identified using DESeq2 (57) with a significance threshold of *P*-adjusted value < 0.05 and fold change ≥ 2. ASEs were analyzed with SUPPA2 (27), and results were filtered based on significance (*P*-adjusted value < 0.05), percent spliced-in (PSI) changes (|ΔPSI| > 0.1), and SUM(TPMs of three replicates) ≥ 2.5. Gene ontology (GO) analysis for functional annotation of candidate alternative spliced transcripts was conducted using the online database for annotation, visualization, and integrated discovery (58).

### Statistical analysis

Statistical analyses were conducted using GraphPad Prism software for experimental data, while data from TCGA and GEO repositories were analyzed using R software. Comparative analyses between experimental groups and their respective controls were performed using Student’s *t*-test, log-rank test, Wilcoxon matched-paired signed-rank test, Mann-Whitney *U* test, and one-way analysis of variance (ANOVA), as appropriate. Data are presented as mean ± SD, and statistical significance was defined as *P* < 0.05.

## Supplemental Material

Supplemental material is available online only. FIG S1A-E, PDF file, 1.6 MB.

FIG S1E-I, PDF file, 1.9 MB.

FIG S2, PDF file, 297 KB.

FIG S3, PDF file, 320 KB.

FIG S4, PDF file, 158 KB.

Table S1, Word file, 19 KB.

Table S2, Word file, 15 KB.

## Acknowledgements

We would like to thank members of Shou-Jiang Gao’s laboratory for technical assistance and helpful discussions. This work was supported by grants from the National Cancer Institute of the National Institutes of Health (CA096512, CA278812, CA284554, CA124332 and CA291244 to S.-J. Gao) and in part by P30CA047904. The authors wish to acknowledge the support of the UPMC Hillman Cancer Center Cytometry Facility (CF), Animal Facility (AF) and Tissue and Research Pathology Services (TARPS), supported by the National Cancer Institute of the National Institutes of Health under award number P30CA047904.

## Author contributions

S. Y. Sun: Investigation, methodology, data curation, formal analysis, visualization, writing–original draft, writing–review. K. Paniagua: Formal analysis, data curation, writing–review. L. Ding: Investigation, data curation, writing–review. X. Wang: Investigation, data curation, writing–review. Y. F. Huang: Formal analysis, writing– review and editing. M. A. Flores: Formal analysis, writing–review and editing. S.-J. Gao: Conceptualization, funding, supervision, managing, formal analysis, investigation, visualization, methodology, writing–original draft, writing–review and editing.

## Data availability

All sequencing data are deposited in GEO. All other data are available upon reasonable request.

