## Supplemental Materials for "KSHV Reprograms Host RNA Splicing via FAM50A to Activate STAT3 and Drive Oncogenic Cellular Transformation"

**Table S1** Summary of the clinical survival data of top nine splicing factors essential for KSHV-induced cellular transformation identified by CRISPR-Cas9 screening in multiple cancer types.

**Table S2** Primers and sequences.

**FIG S1** Clinical survival data of top nine splicing factors essential for KSHV-induced cellular transformation identified by CRISPR-Cas9 screening in multiple cancer types. (A-I) Kaplan-Meier survival curves for the top nine splicing factors in various cancers, including FAM50A (A), NAA38 (B), RBM22 (C), U2AF2 (D), PRMT5 (E), MAGOH (F), RBMX2 (G), SNRPB (H), and PRPF40A (I). Survival outcomes were analyzed based on gene expression levels, with statistical significance determined by log-rank tests.

**FIG S2** (A) Tumor growth trajectories of individual tumors in nude mice implanted with FAM50A knockout KMM cells (sgFAM50A-1, 2 and 3), no sgRNA KMM-Cas9 cells, and control KMM-Cas9 cells with scrambled sgRNAs. (B) Tumor volumes at the study endpoint for each group. Statistical comparisons were performed between each sgRNA-treated group and the Control. Data are presented as mean ± 95% confidence interval (CI), with significance determined using Student's *t*-test. (****$P < 0.0001$).

**FIG S3** Alternative splicing analysis in FAM50A knockout cells. (A-B) Heatmaps displaying $\Delta$PSI values for all differential spliced transcripts between MM vs. FAM50A knockout MM cells (A), and KMM vs. FAM50A knockout KMM cells (B). ΔPSI values were calculated based on three independent biological replicates per group.

**FIG S4** Quantification of SHP2 isoform ratios. (A) SHP2-S/SHP2-L ratio derived from the data in Figure 5A. (B) SHP2-S/SHP2-L ratio calculated from the data in Figure 5C.

**Table S1** Summary of the clinical survival data of top nine splicing factors essential for KSHV-induced cellular transformation identified by Crispr-Cas9 screening in multiple cancer types

| | NAA38 | RBM22 | U2AF2 | PRMT5 | MAGOH | FAM50A | RBMX2 | SNRPB | PRPF40A | Case number |
|---|---|---|---|---|---|---|---|---|---|---|
| LIHC | 0.00072 | 0.0031 | 0.0001 | 0.00022 | 0.083 | 0.026 | 0.0014 | 0.00013 | 0.079 | 365 |
| ACC | | | 0.056 | | 0.00015 | 0.0014 | | | 0.0019 | 79 |
| BLCA | | | | 0.028 | | | | | | 406 |
| COAD | 0.067 | | | | | | | 0.021 | | 279 |
| ESCA | | | | | | 0.052 | 0.0043 | | | 184 |
| HNSC | | | | 0.026 | | | 0.0036 | | | 519 |
| KICH | | 0.013 | | | | 0.016 | | | 0.084 | 65 |
| KIRC | | | | | | 0.0047 | 0.057 | 0.004 | | 531 |
| KIRP | | 0.0071 | | | 0.0045 | | 0.072 | 0.0001 | 0.012 | 287 |
| LAML | | | 0.012 | | | 0.0018 | | 0.047 | | 163 |
| LGG | | | 0.01 | | 0.0001 | | | 0.0004 | 0.0019 | 511 |
| LUAD | | | | | | | | | 0.029 | 502 |
| LUSC | | | | | | | | 0.086 | | 494 |
| MESO | 0.028 | | 0.0002 | | 0.00016 | 0.0016 | | 0.026 | 0.0056 | 85 |
| PAAD | | | | | | | | | 0.0003 | 177 |
| SARC | | 0.056 | 0.012 | | 0.011 | 0.072 | | 0.083 | | 259 |
| THCA | | | | 0.078 | | | | | | 504 |
| UCEC | | | | | | | 0.015 | | 0.097 | 534 |
| UVM | | | 0.038 | | | 0.0019 | | 0.013 | | 80 |

*P*-values are based on comparison between the top 25% and the lower 75% cases.

**Table S2** Primers and sequences

| Primer name | Sequence |
|---|---|
| FAM50A_Rat_Forward | GAAGCAGAGGATTGCAGAGG |
| FAM50A_Rat_Reverse | TTTCTCGCTCCTTCACCAGT |
| Rat_SHP2_total_Forward | GGTTCCACGGTCACTTGTCT |
| Rat_SHP2_total_Reverse | TGGACTTGCTGTCATTGCTC |
| Rat_SHP2_long_Forward | ACAAGCTCTACTCCAGGGAAAC |
| Rat_SHP2_long_Reverse | CACCTTTCTCTCGGATGATG |
| Rat_SHP2_short_Forward | TCGGACAAGGAAACACAGAG |
| Rat_SHP2_short_Reverse | CGATGTCACAGTCCACACCT |
| Rat_SHP2_AS_validation_Forward | GCGCATGACTACACCTTACG |
| Rat_SHP2_AS_validation_Reverse | CCAGGTCCGAAAGTGGTACT |
| pCDH_Rat_SHP2_OE_Forward | TATGAATTCGCCACCATGGACTACAAAGACGATGACGACAAGATGACATCCCGGAGATGG |
| pCDH_Rat_SHP2_OE_Reverse | ATAGCGGCCGCTCATCTGAAACTCCTCTGCT |

### Figure S1A-E

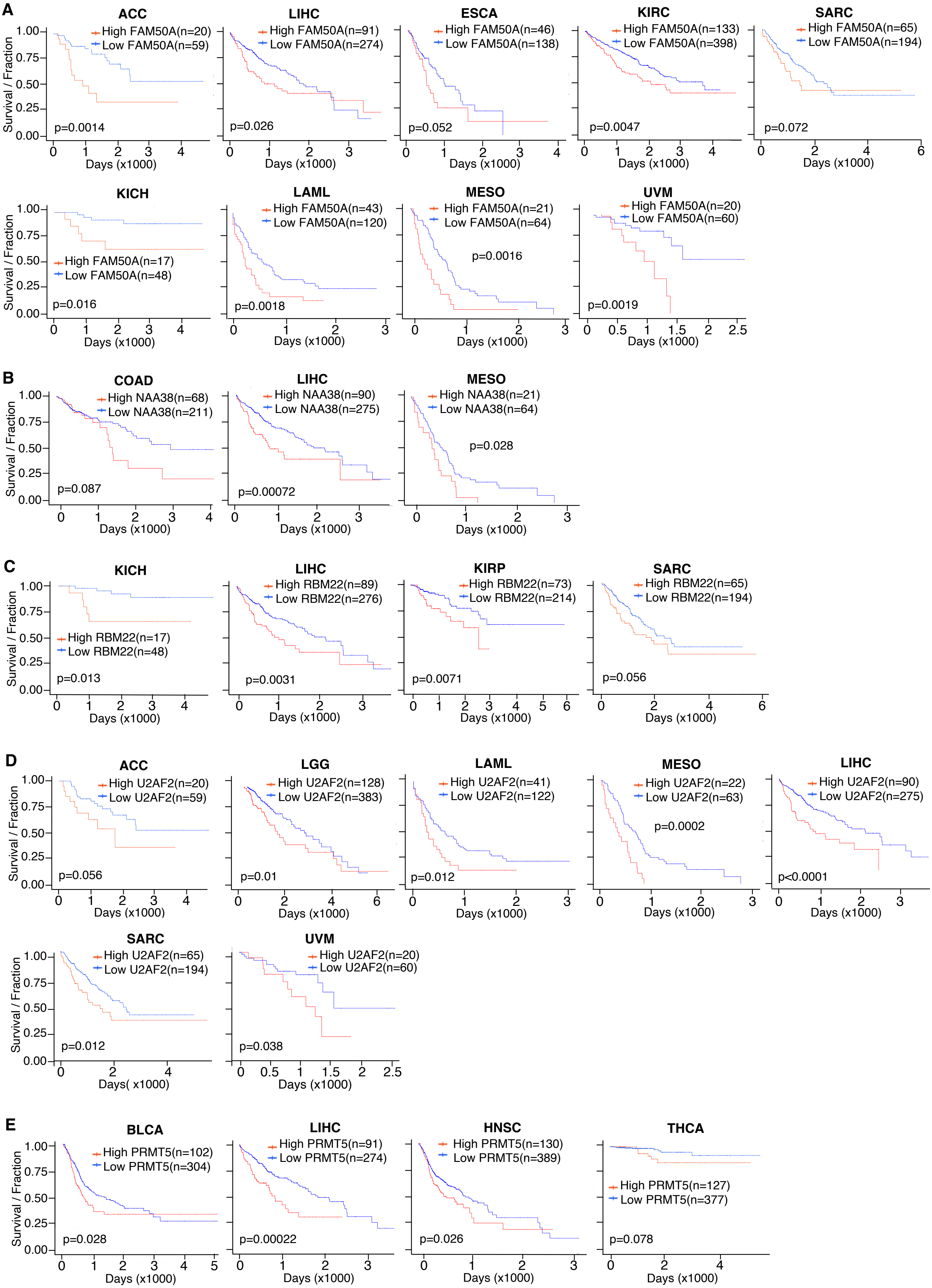
A
ACC
High FAM50A(n=20)
Low FAM50A(n=59)
p=0.0014
LIHC
High FAM50A(n=91)
Low FAM50A(n=274)
p=0.026
ESCA
High FAM50A(n=46)
Low FAM50A(n=138)
p=0.052
KIRC
High FAM50A(n=133)
Low FAM50A(n=398)
p=0.0047
SARC
High FAM50A(n=65)
Low FAM50A(n=194)
p=0.072
KICH
High FAM50A(n=17)
Low FAM50A(n=48)
p=0.016
LAML
High FAM50A(n=43)
Low FAM50A(n=120)
p=0.0018
MESO
High FAM50A(n=21)
Low FAM50A(n=64)
p=0.0016
UVM
High FAM50A(n=20)
Low FAM50A(n=60)
p=0.0019
B
COAD
High NAA38(n=68)
Low NAA38(n=211)
p=0.087
LIHC
High NAA38(n=90)
Low NAA38(n=275)
p=0.00072
MESO
High NAA38(n=21)
Low NAA38(n=64)
p=0.028
C
KICH
High RBM22(n=17)
Low RBM22(n=48)
p=0.013
LIHC
High RBM22(n=89)
Low RBM22(n=276)
p=0.0031
KIRP
High RBM22(n=73)
Low RBM22(n=214)
p=0.0071
SARC
High RBM22(n=65)
Low RBM22(n=194)
p=0.056
D
ACC
High U2AF2(n=20)
Low U2AF2(n=59)
p=0.056
LGG
High U2AF2(n=128)
Low U2AF2(n=383)
p=0.01
LAML
High U2AF2(n=41)
Low U2AF2(n=122)
p=0.012
MESO
High U2AF2(n=22)
Low U2AF2(n=63)
p=0.0002
LIHC
High U2AF2(n=90)
Low U2AF2(n=275)
p<0.0001
SARC
High U2AF2(n=65)
Low U2AF2(n=194)
p=0.012
Days( x1000)
UVM
High U2AF2(n=20)
Low U2AF2(n=60)
p=0.038
E
BLCA
High PRMT5(n=102)
Low PRMT5(n=304)
p=0.028
LIHC
High PRMT5(n=91)
Low PRMT5(n=274)
p=0.00022
HNSC
High PRMT5(n=130)
Low PRMT5(n=389)
p=0.026
THCA
High PRMT5(n=127)
Low PRMT5(n=377)
p=0.078
Survival / Fraction
Days (x1000)

### Figure S1F-I

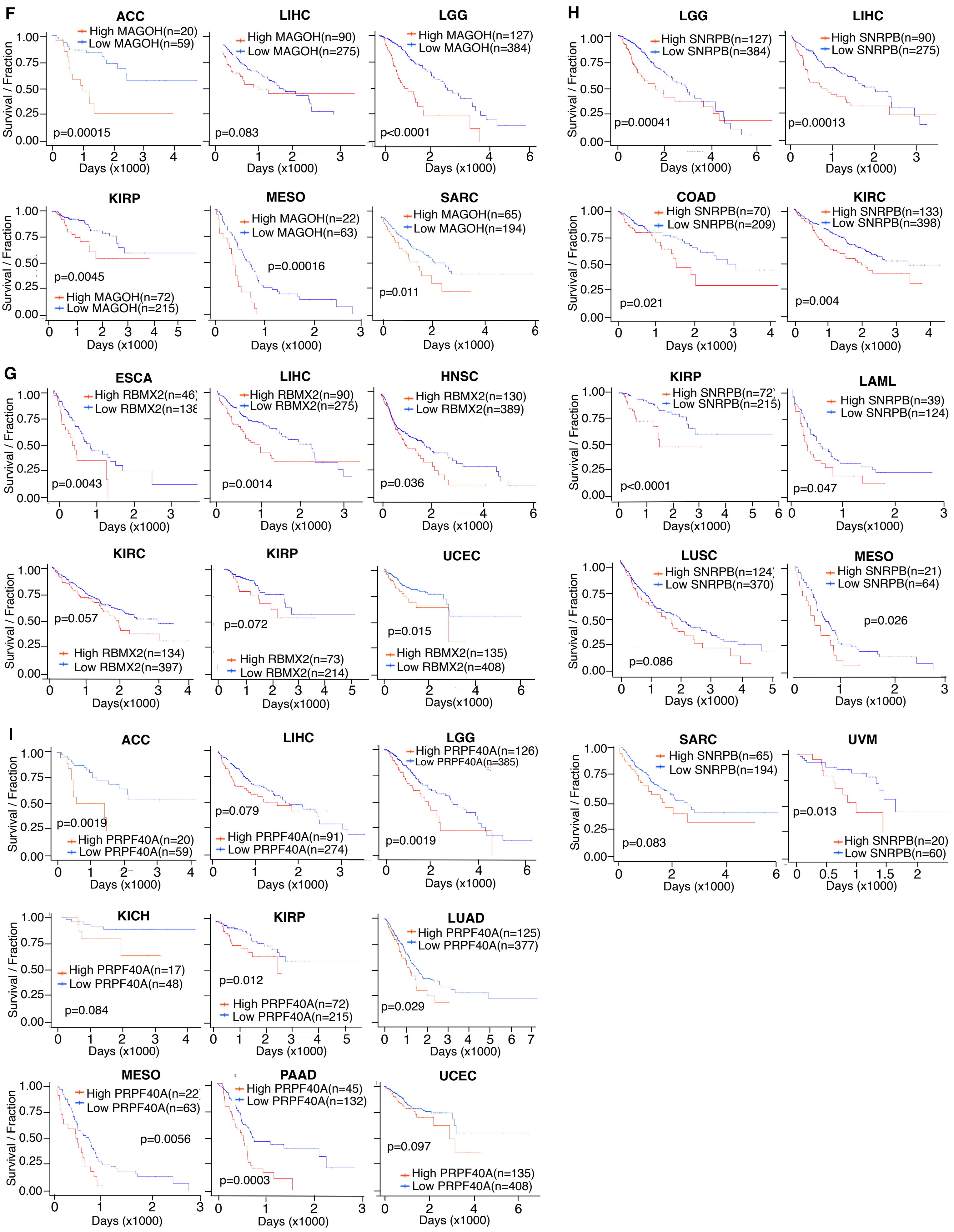

F
ACC
High MAGOH(n=20)
Low MAGOH(n=59)
p=0.00015
LIHC
High MAGOH(n=90)
Low MAGOH(n=275)
p=0.083
LGG
High MAGOH(n=127)
Low MAGOH(n=384)
p<0.0001
KIRP
p=0.0045
High MAGOH(n=72)
Low MAGOH(n=215)
MESO
High MAGOH(n=22)
Low MAGOH(n=63)
p=0.00016
SARC
High MAGOH(n=65)
Low MAGOH(n=194)
p=0.011
H
LGG
High SNRPB(n=127)
Low SNRPB(n=384)
p=0.00041
LIHC
High SNRPB(n=90)
Low SNRPB(n=275)
p=0.00013
COAD
High SNRPB(n=70)
Low SNRPB(n=209)
p=0.021
KIRC
High SNRPB(n=133)
Low SNRPB(n=398)
p=0.004
G
ESCA
High RBMX2(n=46)
Low RBMX2(n=138
p=0.0043
LIHC
High RBMX2(n=90)
Low RBMX2(n=275)
p=0.0014
HNSC
High RBMX2(n=130)
Low RBMX2(n=389)
p=0.036
KIRP
High SNRPB(n=72)
Low SNRPB(n=215)
p<0.0001
LAML
High SNRPB(n=39)
Low SNRPB(n=124)
p=0.047
KIRC
p=0.057
High RBMX2(n=134)
Low RBMX2(n=397)
KIRP
p=0.072
High RBMX2(n=73)
Low RBMX2(n=214)
UCEC
p=0.015
High RBMX2(n=135)
Low RBMX2(n=408)
LUSC
High SNRPB(n=124)
Low SNRPB(n=370)
p=0.086
MESO
High SNRPB(n=21)
Low SNRPB(n=64)
p=0.026
I
ACC
p=0.0019
High PRPF40A(n=20)
Low PRPF40A(n=59)
LIHC
p=0.079
High PRPF40A(n=91)
Low PRPF40A(n=274)
LGG
High PRPF40A(n=126)
Low PRPF40A(n=385)
p=0.0019
SARC
High SNRPB(n=65)
Low SNRPB(n=194)
p=0.083
UVM
p=0.013
High SNRPB(n=20)
Low SNRPB(n=60)
KICH
High PRPF40A(n=17)
Low PRPF40A(n=48)
p=0.084
KIRP
p=0.012
High PRPF40A(n=72)
Low PRPF40A(n=215)
LUAD
High PRPF40A(n=125)
Low PRPF40A(n=377)
p=0.029
MESO
High PRPF40A(n=22)
Low PRPF40A(n=63)
p=0.0056
PAAD
High PRPF40A(n=45)
Low PRPF40A(n=132)
p=0.0003
UCEC
p=0.097
High PRPF40A(n=135)
Low PRPF40A(n=408)
Survival / Fraction
Days (x1000)

#### Figure S2

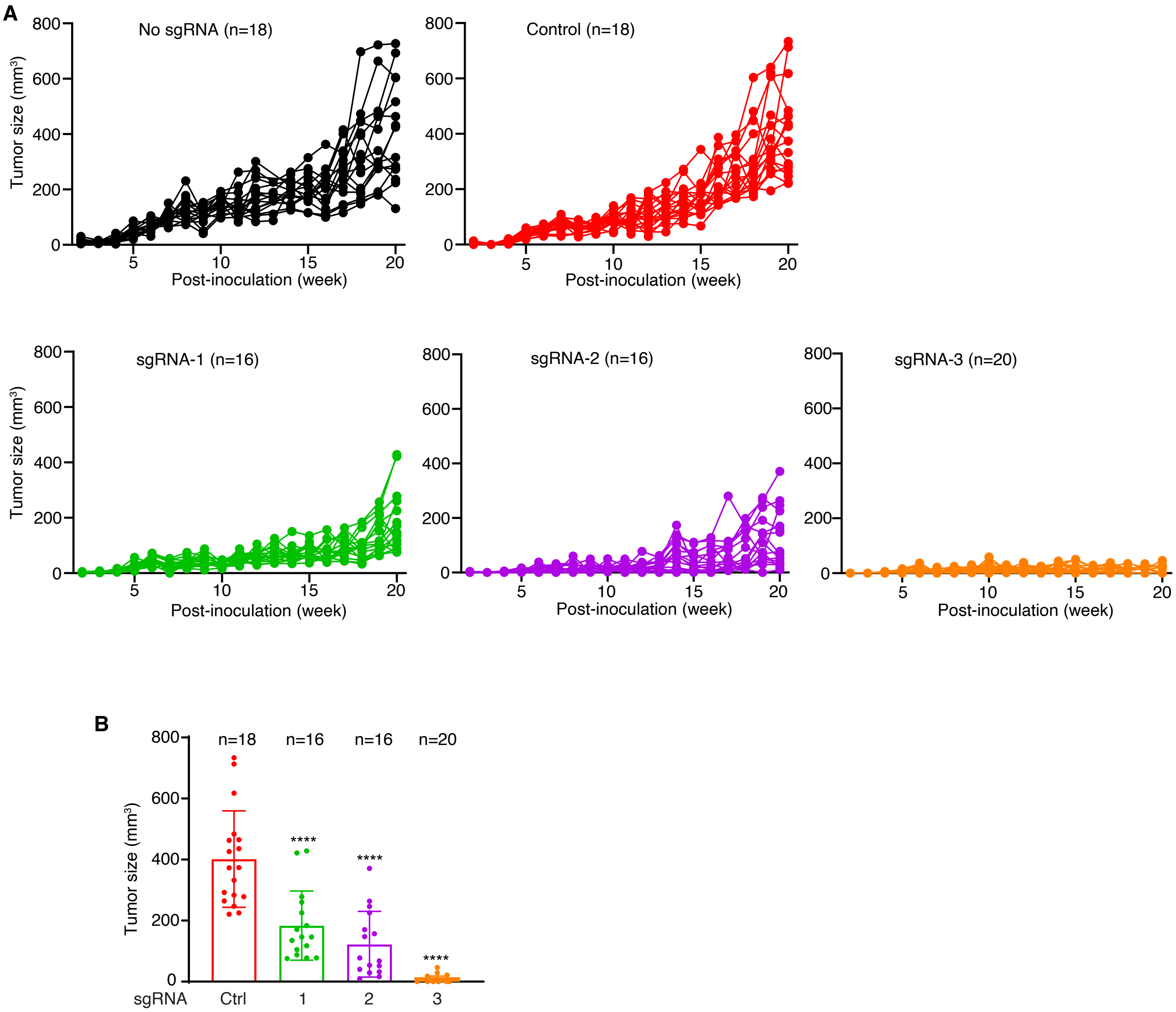

A
No sgRNA (n=18)
Control (n=18)
sgRNA-1 (n=16)
sgRNA-2 (n=16)
sgRNA-3 (n=20)
Tumor size (mm³)
Post-inoculation (week)
0
200
400
600
800
5
10
15
20
B
n=18
n=16
n=16
n=20
****
****
****
sgRNA
Ctrl
1
2
3

### Figure S3

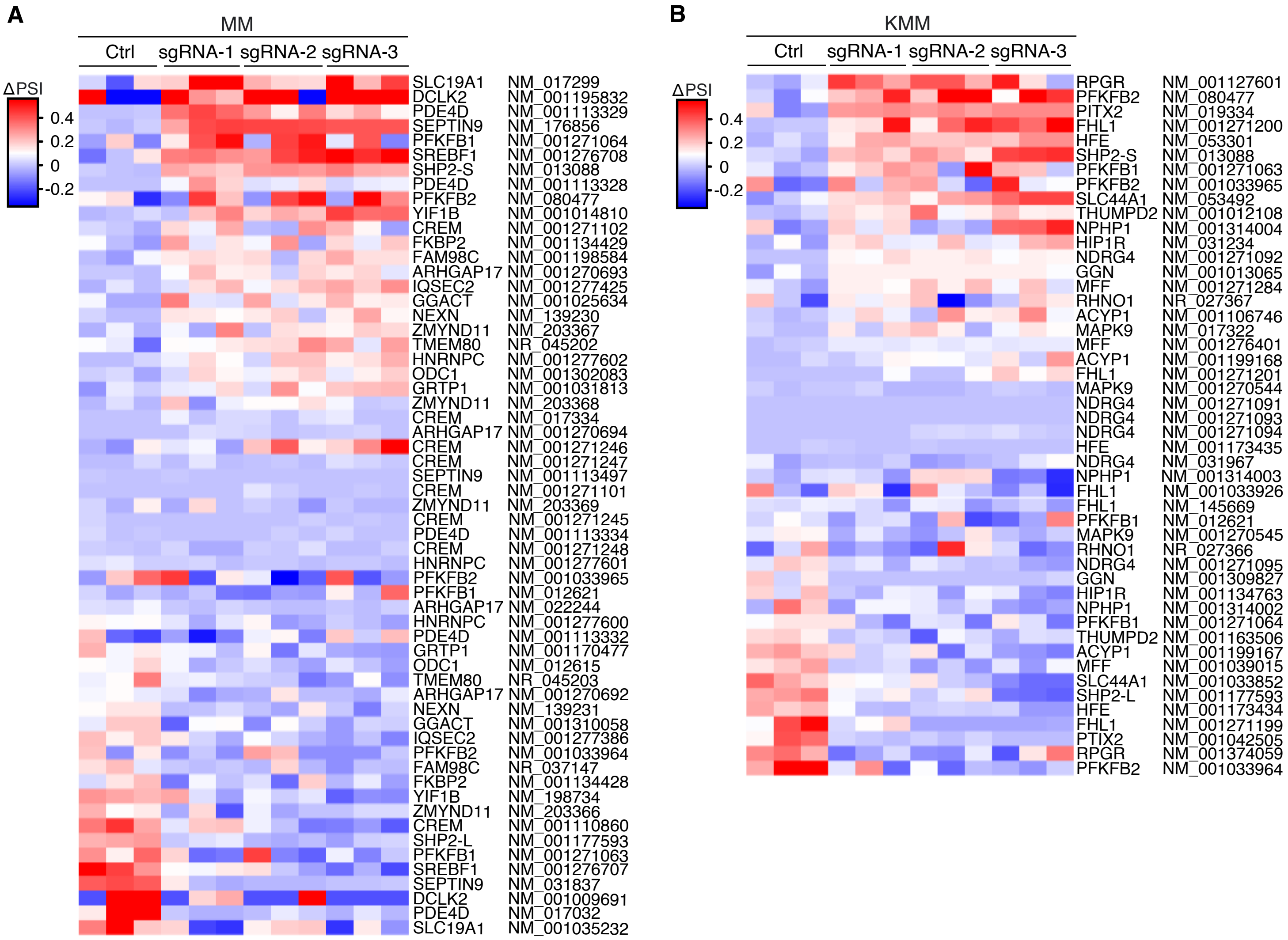
A
MM
Ctrl
sgRNA-1
sgRNA-2
sgRNA-3
ΔPSI
0.4
0.2
0
-0.2
SLC19A1 NM_017299
DCLK2 NM_001195832
PDE4D NM_001113329
SEPTIN9 NM_176856
PFKFB1 NM_001271064
SREBF1 NM_001276708
SHP2-S NM_013088
PDE4D NM_001113328
PFKFB2 NM_080477
YIF1B NM_001014810
CREM NM_001271102
FKBP2 NM_001134429
FAM98C NM_001198584
ARHGAP17 NM_001270693
IQSEC2 NM_001277425
GGACT NM_001025634
NEXN NM_139230
ZMYND11 NM_203367
TMEM80 NR_045202
HNRNPC NM_001277602
ODC1 NM_001302083
GRTP1 NM_001031813
ZMYND11 NM_203368
CREM NM_017334
ARHGAP17 NM_001270694
CREM NM_001271246
CREM NM_001271247
SEPTIN9 NM_001113497
CREM NM_001271101
ZMYND11 NM_203369
CREM NM_001271245
PDE4D NM_001113334
CREM NM_001271248
HNRNPC NM_001277601
PFKFB2 NM_001033965
PFKFB1 NM_012621
ARHGAP17 NM_022244
HNRNPC NM_001277600
PDE4D NM_001113332
GRTP1 NM_001170477
ODC1 NM_012615
TMEM80 NR_045203
ARHGAP17 NM_001270692
NEXN NM_139231
GGACT NM_001310058
IQSEC2 NM_001277386
PFKFB2 NM_001033964
FAM98C NR_037147
FKBP2 NM_001134428
YIF1B NM_198734
ZMYND11 NM_203366
CREM NM_001110860
SHP2-L NM_001177593
PFKFB1 NM_001271063
SREBF1 NM_001276707
SEPTIN9 NM_031837
DCLK2 NM_001009691
PDE4D NM_017032
SLC19A1 NM_001035232
B
KMM
Ctrl
sgRNA-1
sgRNA-2
sgRNA-3
ΔPSI
0.4
0.2
0
-0.2
RPGR NM_001127601
PFKFB2 NM_080477
PITX2 NM_019334
FHL1 NM_001271200
HFE NM_053301
SHP2-S NM_013088
PFKFB1 NM_001271063
PFKFB2 NM_001033965
SLC44A1 NM_053492
THUMPD2 NM_001012108
NPHP1 NM_001314004
HIP1R NM_031234
NDRG4 NM_001271092
GGN NM_001013065
MFF NM_001271284
RHNO1 NR_027367
ACYP1 NM_001106746
MAPK9 NM_017322
MFF NM_001276401
ACYP1 NM_001199168
FHL1 NM_001271201
MAPK9 NM_001270544
NDRG4 NM_001271091
NDRG4 NM_001271093
NDRG4 NM_001271094
HFE NM_001173435
NDRG4 NM_031967
NPHP1 NM_001314003
FHL1 NM_001033926
FHL1 NM_145669
PFKFB1 NM_012621
MAPK9 NM_001270545
RHNO1 NR_027366
NDRG4 NM_001271095
GGN NM_001309827
HIP1R NM_001134763
NPHP1 NM_001314002
PFKFB1 NM_001271064
THUMPD2 NM_001163506
ACYP1 NM_001199167
MFF NM_001039015
SLC44A1 NM_001033852
SHP2-L NM_001177593
HFE NM_001173434
FHL1 NM_001271199
PTIX2 NM_001042505
RPGR NM_001374059
PFKFB2 NM_001033964

#### Figure S4

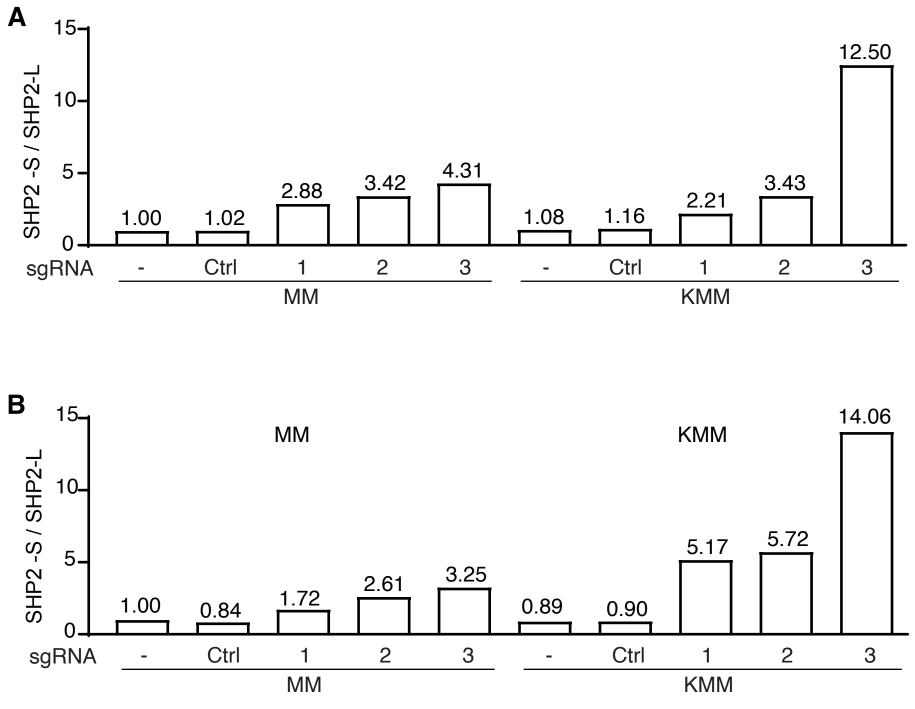
A
15
10
5
0
SHP2 -S / SHP2-L
12.50
1.00
1.02
2.88
3.42
4.31
1.08
1.16
2.21
3.43
sgRNA
-
Ctrl
1
2
3
-
Ctrl
1
2
3
MM
KMM
B
15
10
5
0
SHP2 -S / SHP2-L
MM
KMM
14.06
5.17
5.72
1.00
0.84
1.72
2.61
3.25
0.89
0.90
sgRNA
-
Ctrl
1
2
3
-
Ctrl
1
2
3
MM
KMM
